# Warming can destabilise predator-prey interactions by shifting the functional response from Type III to Type II

**DOI:** 10.1101/498030

**Authors:** Uriah Daugaard, Owen L. Petchey, Frank Pennekamp

**Author notes:** **Corresponding author:** Frank Pennekamp, Institute of Evolutionary Biology and Environmental Studies, University of Zurich, Winterthurerstrasse 190, 8057 Zurich, Switzerland.

## Abstract

1. The potential for climate change and temperature shifts to affect community stability remains relatively unknown. One mechanism by which temperature may affect stability is by altering trophic interactions. The functional response quantifies the *per capita* resource consumption by the consumer as a function of resource abundance and is a suitable framework for the description of nonlinear trophic interactions.
2. We studied the effect of temperature on a ciliate predator-prey pair (*Spathidium sp*. and *Dexiostoma campylum*) by estimating warming effects on the functional response and on the associated conversion efficiency of the predator.
3. We recorded prey and predator dynamics over 24 hours and at three temperature levels (15, 20 and 25°C). To these data we fitted a population dynamic model including the predator functional response, such that the functional response parameters (space clearance rate, handling time, and density dependence of space clearance rate) were estimated for each temperature separately. To evaluate the ecological significance of temperature effects on the functional response parameters we simulated predator-prey population dynamics. We considered the predator-prey system to be destabilised, if the prey was driven extinct by the predator.
4. Effects of increased temperature included a transition of the functional response from a Type III to a Type II and an increase of the conversion efficiency of the predator. The simulated population dynamics showed a destabilisation of the system with warming, with greater risk of prey extinction at higher temperatures likely caused by the transition from a Type III to a Type II functional response.
5. Warming-induced shifts from a Type III to II are not commonly considered in modelling studies that investigate how population dynamics respond to warming. Future studies should investigate the mechanism and generality of the effect we observed and simulate temperature effects in complex food webs including shifts in the type of the functional response as well as consider the possibility of a temperature dependent conversion efficiency.

## Introduction

Temperature is a prime driver of biological systems through the temperature-dependence of biological rates (Brown, Gillooly, Allen, Savage, & West, 2004). Effects of temperature on biological rates of individuals (e.g., metabolic rates) are expected to scale up to the population level and consequently affect carrying capacities and population growth rates (Bernhardt, Sunday, & O’Connor, 2018; Gillooly, Brown, West, Savage, & Charnov, 2001; Savage, Gilloly, Brown, & Charnov, 2004) as well as ecosystem properties such as respiration (Yvon-Durocher, Jones, Trimmer, Woodward, & Montoya, 2010). A major outstanding challenge is to understand how temperature affects species interactions and changes in community structure, dynamics and stability (Burnside, Erhardt, Hammond, & Brown, 2014; Fussmann, Schwarzmüller, Brose, Jousset, & Rall, 2014; Walther, 2010).

Trophic interactions play an important role in ecosystems and have a close link to temperature via the consumption and metabolism of predators and prey (Rall et al., 2012). Food webs have been one focus of empirical and theoretical investigations to understand the effect of temperature on communities and ecosystems (e.g. Doney et al., 2012; Petchey, McPhearson, Casey, & Morin, 1999). Many of these studies show that warming can have profound consequences for the stability of food webs (Fussmann et al., 2014; Rall, Vucic-Pestic, Ehnes, Emmerson, & Brose, 2010; Uszko, Diehl, Englund, & Amarasekare, 2017; Vasseur & McCann, 2005). Nevertheless, there are gaps in our understanding about the mechanisms by which temperature affects trophic interactions.

The functional response provides a general and succinct conceptualisation of a trophic interaction. It describes the *per capita* prey consumption rate by the predator as a function of prey abundance (Solomon, 1949). The most commonly used models of the functional response are the disc equations developed by Holling (1959) or variations of them (for an overview see Jeschke, Kopp, & Tollrian, 2002). The main parameters in these models are the space clearance rate *a* (units: area or volume per time; often referred to as attack rate or search rate) and handling time *h* (unit: time). Real (1977) generalised the functional response by introducing the possibility of distinguishing between a resource dependent and a resource independent space clearance rate *a* = *bN^q^*. Here, *N* is the resource abundance, *b* is a constant and *q* is the space clearance rate exponent (also known as scaling exponent and attack exponent or as its transferable version, the Hill exponent H, with *H* = *q* + 1). To simplify notation, henceforth we will refer to space clearance rate exponent as the exponent *q*. This definition of the space clearance rate allows classification of the functional response into a Type II (*q* = 0, meaning resource independent space clearance rates *a* = *b*) or a Type III (*q* > 0, i.e. space clearance rate increases with prey abundance). The Type III response is generally considered to be stabilising as there is less predation pressure at low prey densities (e.g. Uszko, Diehl, Pitsch, Lengfellner, & Müller, 2015; Yodzis & Innes, 1992). The exponent *q* is therefore a key parameter influencing the stability properties of a predator-prey system.

How functional response parameters vary with traits or abiotic conditions has been the focus of many studies (Jeschke et al., 2002; Kalinoski & DeLong, 2016; Pritchard, Paterson, Bovy, & Barrios-O’Neill, 2017). The temperature dependence of handling time and of the resource independent space clearance rate has repeatedly been studied (e.g. Sentis, Hemptinne, & Brodeur, 2012; Thompson, 1978; Uiterwaal & DeLong, 2018; Vucic-Pestic, Ehnes, Rall, & Brose, 2011; Zamani, Talebi, Fathipour, & Baniameri, 2006). The metabolic theory of ecology states that the metabolic rate scales exponentially with temperature (MTE, Brown et al., 2004). However, two meta-analyses concluded that the MTE is generally not suited for predicting the temperature dependence of the handling time *h* and of the resource independent space clearance rate *a* (Englund, Öhlund, Hein, & Diehl, 2011; Rall et al., 2012). Hence, there is still uncertainty surrounding the general influence of temperature on these parameters.

Little is known about how the exponent *q* varies with temperature. Although some studies have investigated the relationship of *q* with body mass and habitat structure (Barrios-O’Neill et al., 2016; Kalinkat et al., 2013), we currently lack studies exploring the temperature dependence of *q* (but see Uszko et al., 2017). Further, similar to the exponent *q* the temperature dependence of the predator’s conversion efficiency is insufficiently understood. The conversion efficiency is the number of predators produced per prey consumed. Modelling studies (e.g. Fussmann et al., 2014; Uszko et al., 2017; Vasseur & McCann, 2005) tend to assume temperature independent conversion efficiency and a fixed Type II or III functional response across temperature, ignoring implications of these choices for the stability of the system.

Despite the unknown relationship between *q* and temperature, temperature driven type shifts of the functional response have been reported before. In fact, both the stabilising (from Type II to Type III, e.g. Mohaghegh, De Clercq, & Tirry, 2001; South & Dick, 2017; Wang & Ferro, 1998; Ziaei Madbouni, Samih, Namvar, & Biondi, 2017) and the destabilising transition (from Type III to Type II, see Dong, Liu, Xie, Cong, & Wang, 2017; Taylor & Collie, 2003) have previously been found. In some cases, the functional response shifted back and forth between types with warming (Eggleston, 1990; Mondal, Chandra, Bandyopadhyay, & Ghosh, 2017). These shifts have been found predominantly in insect consumer-resource pairs and sporadically in crustacean-molluscs and fish-crustacean pairs. However, to our knowledge, these shifts have never been observed through the direct estimation of *q*. Rather, studies most often used the method described by Juliano (2001) or similar categorical approaches, in which the functional response type is limited to a Type II (i.e. *q* = 0) or a classical Type III (i.e. *q* = 1) and is determined for instance with a logistic regression of a polynomial function. An exception is the Taylor and Collie (2003) study, which used a linear regression approach to determine the functional response type, but which neglected the prey depletion present in the functional response experiment (Rogers, 1972). It has been pointed out recently that categorical approaches to determine the functional response type such as the Juliano (2001) method might be inadequate to correctly evaluate subtle changes at low prey densities (Barrios-O’Neill, Dick, Emmerson, Ricciardi, & MacIsaac, 2015). Hence, the validity of the above mentioned type transitions needs to be addressed with appropriate methodology. Furthermore, none of the studies that reported type shifts analysed the dynamic consequences for the stability of the studied consumer-resource pair.

Ciliates are convenient study organisms for many ecological and evolutionary questions (see Altermatt et al., 2015). However, one of their convenient features, i.e. their short generation times, makes the estimation of functional response parameters from feeding trials challenging. Changes in prey abundance may not only result from consumption, but also prey reproduction, which complicates parameter estimation. A recently developed maximum likelihood method makes it possible to take the growth and mortality rate of the prey into account for the estimation of parameters (Rosenbaum & Rall, 2018). We used this method to investigate the temperature dependence of the functional response, including the possibility of type-shifts via the exponent *q* and of varying conversion efficiencies *c*. We used a microbial predator-prey system consisting of the predatory ciliate *Spathidium sp*. and its ciliate prey *Dexiostoma campylum*. The null hypotheses were that temperature has no effect on any of the parameters. Since we found evidence against these hypotheses, we assessed the ecological significance of the observed temperature-dependencies of these parameters on the stability of the predator-prey interaction. We classified the predator-prey system as destabilised if the prey was driven extinct by the predator.

## Materials and Methods

### Data acquisition: functional response experiments

We carried out functional response experiments with *Spathidium sp*. as the predator and *Dexiostoma campylum* as the prey, which co-occur in nature (e.g. Biyu, 2000). The experiments were done at temperatures of 15 °C, 20 °C and 25 °C. Both species were kept at their respective treatment temperatures prior to the experiment for an acclimatisation period of six weeks. We maintained *D. campylum* in a bacterised organic protozoan pellet medium (Carolina Biological Supply Company, Burlington NC; concentration of 0.55 gL^−1^, see Altermatt et al., 2015). For the functional response experiment, we increased the concentration to 1.1 gL^−1^ to achieve the high prey densities required for the experiment. Predator individuals were kept in three 12-well plates and fed *ad libitum* with *D. campylum* and *Colpidium striatum*.

For the experiment, we used eight prey density levels, of which six were the same across temperature (50, 100, 250, 500, 750 and 1000 individuals per millilitre) and two were adjusted to the prey densities reached in the maintenance cultures (1500 & 3075, 1500 & 2890 and 2000 & 4177 individuals per millilitre respectively at 15, 20 and 25 °C). Prey density was manipulated by dilution. For each temperature and density level we had nine replicates, six with predators present (five predator individuals added) and three control replicates (no predators added). We used 24-well plates and on each we distributed two treatment replicates and one control replicate belonging to a temperature level. By using block randomisation, we randomly assigned each well within a plate to a density and treatment / control (predator or no predator).

To improve the comparability of the experiment across the three temperature levels, we pre-fed the predators according to the temperature at which they were maintained. We adjusted the feeding rates and amounts for each temperature such that the starvation period prior the experiment was long enough for digestion but not so long that the individuals would show signs of food shortage. The length of starvation period was determined based on laboratory observations during the acclimatisation period.

Predator individuals were pipetted from the maintenance plates to the wells assigned to the treatment following a randomisation pattern. After the addition, we confirmed that each well contained the correct number of predators and filled each well with 1 mL of prey culture. Plates were incubated for 24 hours in temperature-controlled incubators with positions in the incubators chosen randomly. Evaporation during the 24 hour period was limited by placing sterile jars containing deionised water alongside the plates. Plates were kept in the dark during the experiment.

After the incubation period, we manually counted predators using a light microscope to record predator growth. To estimate the prey densities we homogenised the volume in a well by gently pipetting the liquid three times. We then sampled 0.6 mL from each well by pipetting it into a counting chamber and took three consecutive 5 seconds videos of different (non-overlapping) regions of the chamber, which together covered approximately 50 % (0.5 mL) of each well. Videog-raphy involved a Hamamatsu C11440 camera, a Leica M205 C dissecting microscope with dark field illumination, and the software HCImage Live. We used the R package bemovi (Pennekamp, Schtickzelle, & Petchey, 2015) to estimate prey density.

### Constructing the model: accounting for predator and prey growth and mortality in the functional response estimation

Predation can be modelled with the generalised functional response equation derived by Real (1977, equation 1):

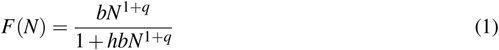

where *N* is the prey abundance and *F* (*N*) is the *per capita* rate of prey consumption by the predator. The parameters are the handling time *h* (unit: time) and the constant *b* (unit: volume per time) and exponent *q* (dimensionless) of the space clearance rate *a* = *bN^q^*. The exponent *q* is related to the Hill exponent *H* (*H* = *q* + 1). The functional response can be a Type I (linear, *q* = 0 and *h* = 0), a Type II (hyperbolic curve, *q* = 0, *h* > 0) or a Type III (sigmoid curve, *q* > 0, *h* > 0).

In functional response experiments, prey abundance usually decreases over time if consumed prey items are not replaced. This is known as prey depletion (Rogers, 1972; Rosenbaum & Rall, 2018). The total prey depletion is *F*(*N*)*TP*, where *T* is duration of the experiment and *P* is the predator abundance. If predation is the only process affecting prey density we expect the instantaneous rate of change in prey density *dN*/*dt* to be equal to the negative of the total prey consumption rate *F*(*N*)*P*:

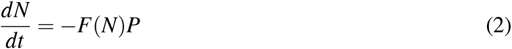

Equation 2 is only correct if the growth and background mortality rates of the prey are negligibly small. In other words, only if the time interval considered is sufficiently short so that there is no or almost no change in prey density caused by other reasons than predation. Following Rosenbaum and Rall (2018), growth and natural mortality of the prey can be taken into account by extending equation 2 with the addition of a logistic growth term (equation 3):

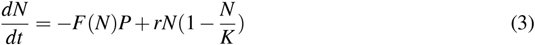

The added parameters are the carrying capacity *K* (unit: individuals per volume) and the intrinsic growth rate *r* (unit: per time). The logistic growth term implies natural prey mortality when *N* > *K* and prey growth when *N* < *K*.

Equations 2 and 3 assume that the predator density is constant. This is only justified if the effects of the predator’s growth and mortality rates are negligible in the time frame considered. Otherwise, the predator’s density change over time can be added to the above model by including the numerical response (Solomon, 1949). The numerical response describes the change in predator abundance as a function of prey abundance (equation 4).

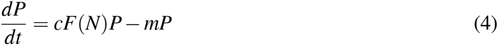

*dP*/*dt* is the rate of change in predator density. The parameter *c* is the conversion efficiency (dimensionless), which determines by how much the total prey consumption rate *F*(*N*)*P* increases the predator density. Also included in equation 4 is the mortality rate *m* of the predator (unit: per time). Together, equations 3 and 4 closely resemble the Rosenzweig-MacArthur model (Rosenzweig & MacArthur, 1963), with the difference that the functional response can be either Type I, II or III.

Starvation or other stressors can induce predator encystment with cysts being able to survive extended periods of time (e.g. Moore, 1924). The detection of the cysts with a light microscope within the experiment wells is relatively straightforward. During the experiment we only encountered predator growth and observed no or close to no cysts (Fig. S1). Predators could in principle die of membrane perforation due to mechanical forces (e.g. due to pipetting). However, as we confirmed the number and integrity of the predators visually after their addition to the experiment wells, we simplified equation 4 by setting the predator mortality rate to 0 (equation 5). In the section “*Model improvement*” in the supplementary material we discuss how to adjust the model if predator mortality is not negligible.

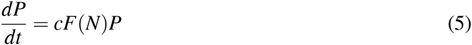

#### Fitting the model

We used the temporal changes in the prey and predator densities recorded in the feeding experiment to investigate the temperature-dependence of the functional response. We applied the method developed by Rosenbaum and Rall (2018), which finds the set of parameter values that provide the best fit to the observed end prey densities. To do so, this method repeatedly and numerically solves equation 3 (prey dynamics) for each given prey density, with different parameter values each time. Each combination of parameter values yields a set of predicted end prey densities. The method calculates the likelihood of each set of predicted end prey densities assuming that each observed end prey density is log-normally distributed around the respective predicted density (with variance estimated by the method). The set of parameter values that results in the predicted end prey densities with the highest likelihood is ultimately chosen as the best fit. In other words, the method is an iterative maximum likelihood method which tests different sets of parameter values and determines which one of them predicts the observed prey density the best.

We extended this method to solve equation 3 along with equation 5 (predator dynamics). We adjusted the maximum likelihood estimation algorithm by adding the likelihood of the predicted end predator abundances. For this, we assumed that the observed end predator abundances follow a Poisson distribution with the predicted end predator abundances as the expected number. Thus, the estimation of the functional response parameters takes into account both the prey and the predator dynamics.

For the numerical solution of the ordinary differential equations (ODEs) we used the package *odeintr* (Keitt, 2017) and for the maximum likelihood estimation we used the function *mle2* from the package *bbmle* (Bolker & Team, 2017). We fitted the model with parameters on the natural log scale to improve convergence, considering that the parameter values can differ by several orders of magnitudes. Subsequently, we refitted the model on the normal scale using the back-transformed solutions obtained on the log-scale to estimate the parameter values and their standard errors on the normal scale. The scale used for the fitting only impacts the convergence stability of the estimation and has no effect on the optimal solution (see the manual of Rosenbaum & Rall, 2018). However, we kept the carrying capacity and the constant *b* on the log-scale, since their magnitudes were too different.

To test whether the inclusion of predator growth significantly improved our model, we compared the AIC values of the models with and without predator growth (i.e. setting the conversion efficiency *c* to 0 for the latter). The model including predator growth showed the better fit across all temperature levels (dAIC values respectively for the comparison at 15, 20 and 25 °C: 1.3, 12.4 and 162.2). Similarly, we also tested whether imposing a classical Type II functional response (i.e. exponent *q* fixed to 0) improved the fit. We found that the model with free *q* showed the better fit at all temperature levels (dAIC of 44.6, 83.3 and 14.88 for temperatures 15, 20 and 25 °C, respectively). Refer to Table S1 for more information about the model comparisons.

We simulated the 95% confidence intervals for the functional response across temperature by first drawing the model parameters 1,000 times randomly from a multivariate normal distribution (mean = estimated parameter values, variance = covariance matrix of model fit). The correlation matrices of the model fits are reported in Table S5 in the supplementary material. We excluded parameter combinations that included biologically meaningless values (i.e. handling times below 0). We then used these parameter combinations to simulate the prey consumption curves. From these we selected the 2.5% and the 97.5% quantiles for our confidence intervals.

#### Stability analysis

To explore the stability implications of the functional response parameter combinations across temperature, we simulated the predator-prey population dynamics. This approach takes into account that temperature simultaneously affects multiple parameters (with potentially counteracting effects) and hence evaluates the dynamic consequences of changes for the whole system rather than testing whether the differences between the specific parameters deviate from zero effect in isolation.

For each temperature level we used 10,000 parameter combinations obtained by sampling randomly from a multivariate normal distribution (mean = estimated parameter values, variance = covariance matrix of model fit), again excluding parameter combinations that contained biologically meaningless values. We used the remaining parameter combinations and equations 3 and 5 to simulate the population dynamics at each temperature for 100 days with time-step *dt* = 0.01d.

As we did not measure mortality experimentally, we collected mortality rates from the literature and scaled rates according to the Metabolic Theory of Ecology (Brown et al., 2004) and equation 6, which is reasonably well supported for the mortality rate (McCoy & Gillooly, 2008; Uszko et al., 2017).

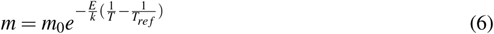

In the equation, *m* is the mortality rate, *m*_0_ is a scaling constant, *E* is the activation energy, *k* is the Boltzmann’s constant (8.62 × 10^−5^ eVK^−1^), *T* is the environmental temperature and *T_ref_* is a reference temperature (in our case 288.15 K, i.e. 15° C).

Mortality rates of various protists species were extracted from Figure 3 of DeLong et al. (2015). Mortality rates ranged approximately from 0.03 to 2.5 d^−1^ (mean = 0.52 d^−1^, sd = 0.58 d^−1^, median = 0.31 d^−1^). Based on this wide range and on the observations during the acclimatisation period, we parameterised the simulations with a predator mortality of 0.1 d^−1^ at 15 °C. We extrapolated the mortality rates at the other temperatures according to equation 6, assuming constant predator mass across the considered temperature range and an activation energy of 0.65 eV (McCoy & Gillooly, 2008). Thus, at 20 °C and at 25 °C the mortality rates were 0.156 d^−1^ and 0.241 d^−1^, respectively. To explore the effect of the predator mortality rates on the population dynamics, we carried out a sensitivity analysis, varying the death rate values. Additionally, to provide a strong argument that changes in the functional response parameters *h*, *b* and *q* drive the destabilization of the system at high temperature, we repeated the simulations with fixed values for the remaining parameters: *r*, *K*, *c* and the estimated standard deviation σ of the log-normal distribution were fixed to the estimated values at 20 °C.

We quantified stability as the percentages of replicate simulations with prey persistence, where the extinction threshold is 1 individual per millilitre. For the cases in which the prey went extinct, we calculated the time to extinction.

## Results

### Estimating the functional response across temperatures

Temperature had a clear effect on changes in prey density under predation and control treatments (Fig. 1). At low and high prey density, prey consumption was visible, whereas at intermediate prey densities the prey growth exceeded consumption, resulting in a hump-shaped pattern of prey change which was consistent across temperatures (Fig. 1A-C). For the temperatures 15 and 20 °C the highest prey densities exceeded the respective estimated carrying capacities (Fig. 2), hence natural prey mortality is expected to occur. As we explicitly model density-dependent growth of the prey, growth (*N* < *K*) and natural mortality (*N* > *K*) are appropriately accounted for. Temperature also influenced predator density, most noticeably when prey density and temperature were high (Fig. S1). The model accurately captured both the prey and the predator dynamics (Fig. 1A-C and Fig. S1)

**Figure 1:**
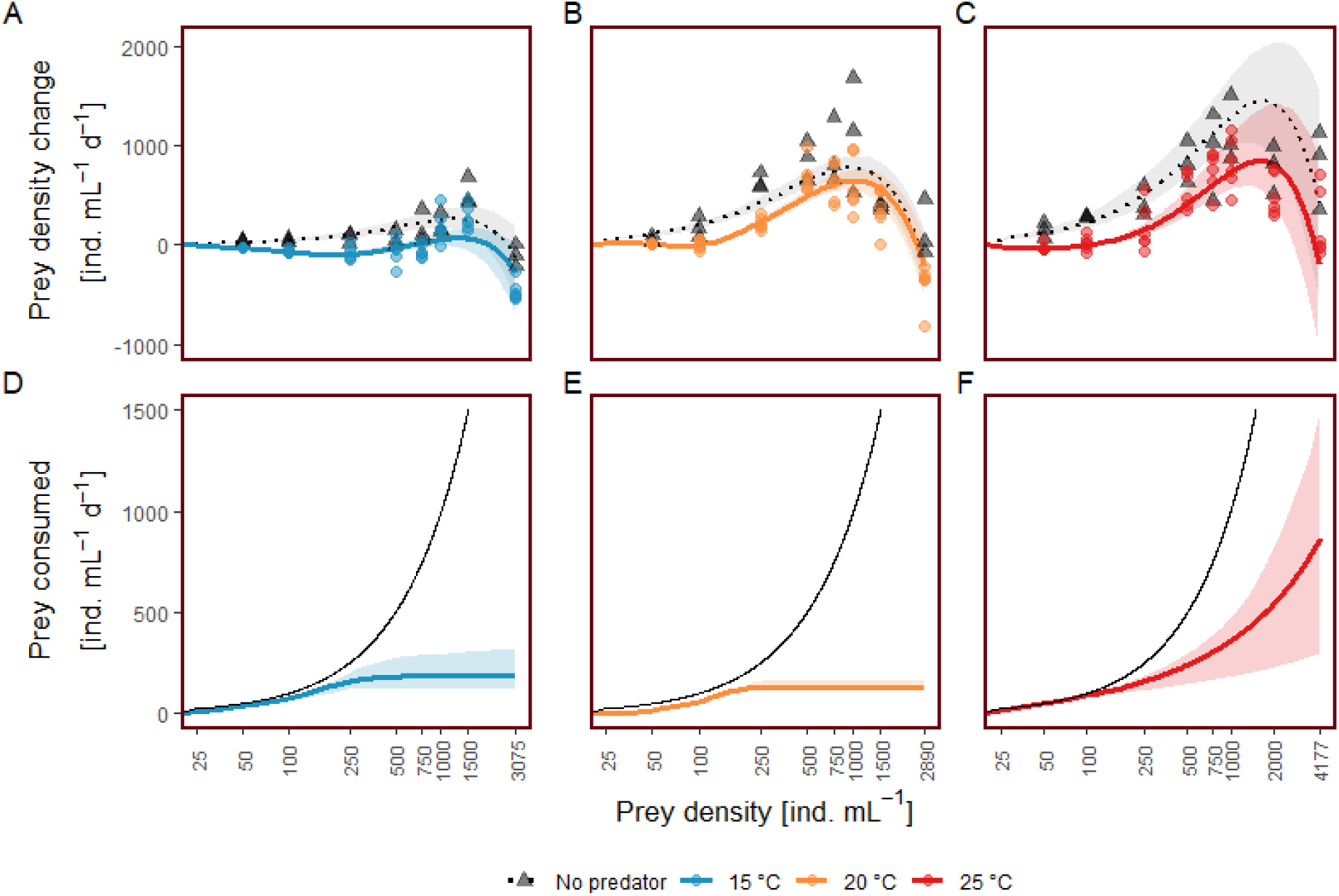
Model fits (lines) and corresponding 95% confidence intervals on the linear-log10 scale. **A) - C)** Functional response across temperature including prey growth (circles are in the presence of predators, triangles in the absence of predators). Negative values on the y-axis denote a reduction in prey density (i.e. prey consumption) and positive values denote an increase (i.e. prey growth). The lines and areas are the fit and confidence intervals. **D) - F)** Daily prey consumption based on functional response parameters (accounting for prey growth or mortality). For the estimation we set the prey growth rate *r* = 0. The black line represents the theoretical maximal prey consumption.

**Figure 2:**
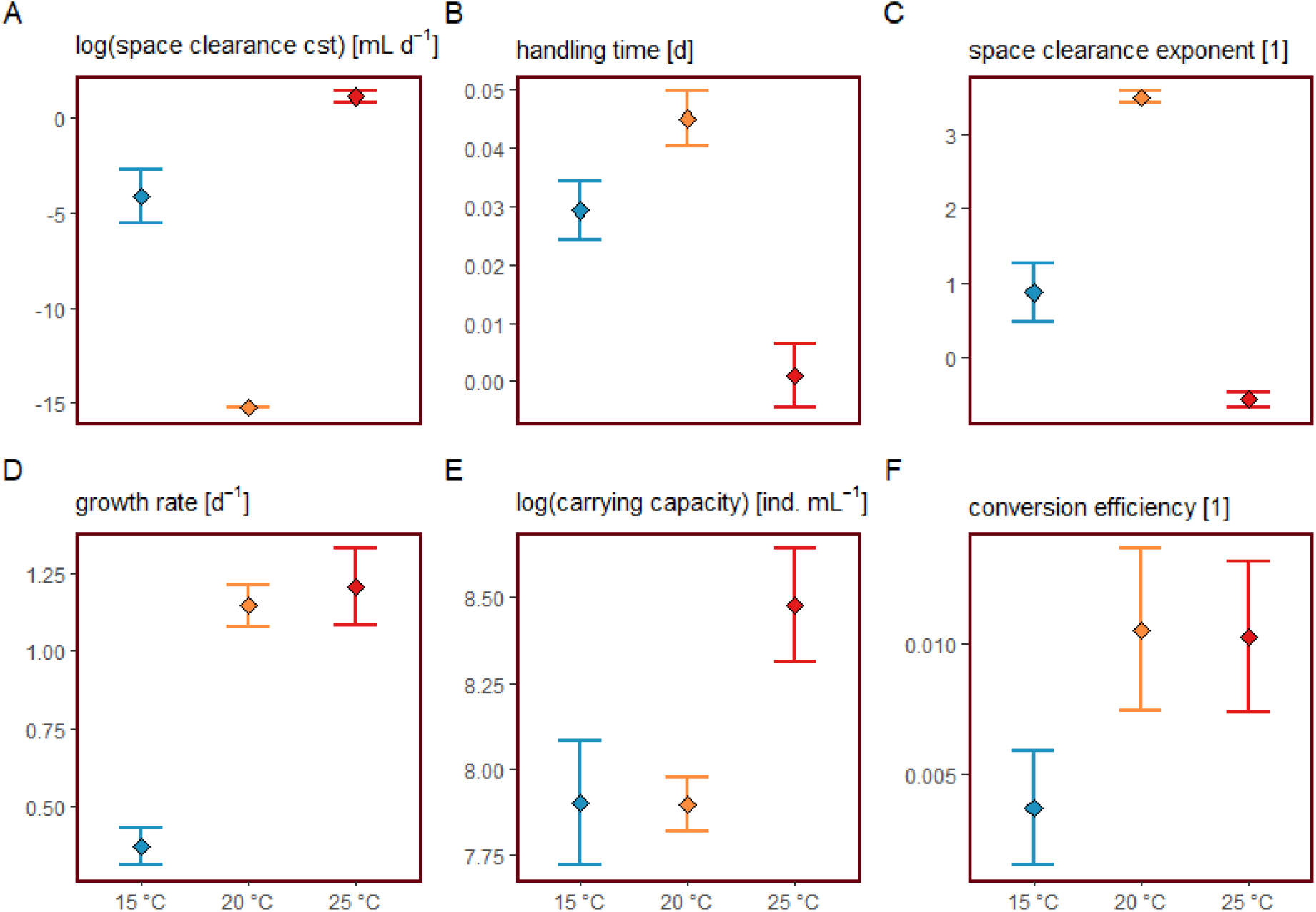
Model parameter estimates (squares) and respective standard errors (bars) across temperature. The respective units are denoted in square brackets and [1] designates a dimensionless parameter **A)** natural log-transformed space clearance rate constant *b*; **B)** handling time *h*; **C)** space clearance rate exponent *q*; **D)** growth rate *r*; **E)** natural log-transformed carrying capacity *K*; **F)** conversion efficiency *c*.

Importantly, the functional response transitioned from a Type III at 15 and 20 °C to a Type II at 25 °C (Fig. 2), and the prey consumption was largest at the highest temperature (Fig. 1D-F, for a close up at low prey density refer to Fig. S2). While the exponent *q* was larger than 0 (i.e. Type III) at 15 and 20 °C, it changed to *q* −0.57 (i.e. Type II) at 25 °C. As Type II is usually associated with *q* = 0, we also fitted the model constraining the exponent to zero (Table S2 in the supplementary material). The results were qualitatively similar and hence the patterns in population dynamics are independent of the approach chosen (Table S4 in the supplementary material).

All of the other functional response parameter values showed temperature dependence (Fig. 2, Table S2). The space clearance constant *b* decreased from 15 to 20 °C, but than increased again at 25 °C. As both the exponent *q* and the constant *b* influence the space clearance rate *a*, they should be interpreted jointly. Hence, Fig. S4 visualises the temperature and prey density dependence of the estimate space clearance rate. Handling time responded in the opposite fashion to the space clearance constant *b*, being similar for 15 and 20 °C but decreased at 25 °C. Consistent with the observed predator growth, the conversion efficiency increased with warming. Further, the growth rate of the prey increased with warming. However, the increase was much bigger between 15 °C and 20 °C (from 0.37 to 1.15) than between 20 °C and 25 °C (from 1.15 to 1.21). Estimated prey carrying capacities were comparable with densities reached in the maintenance jars.

### Population stability

To evaluate the ecological significance of temperature effects on the trophic interaction, we simulated predator-prey population dynamics based on estimated parameters and temperature-dependent mortality rates (*m*_15_ = 0.1d^−1^, *m*_20_ = 0.156d^−1^ and *m*_25_ = 0.241d^−1^, Fig. 3A-B). In the vast majority of simulations, the prey persisted at 15 and 20 °C (Fig. 3C). In contrast, in almost all simulations at 25° C the prey went extinct. Moreover, for cases in which prey went extinct, the time to extinction was much shorter at 25 °C (Fig. 3D). These patterns persisted (although attenuated) across considerably higher mortality rates as well as in simulations where we prevented negative values for the exponent *q* and as well as in those where we fixed all parameters except *h*, *b* and *q* (see Tables S3-4).

**Figure 3:**
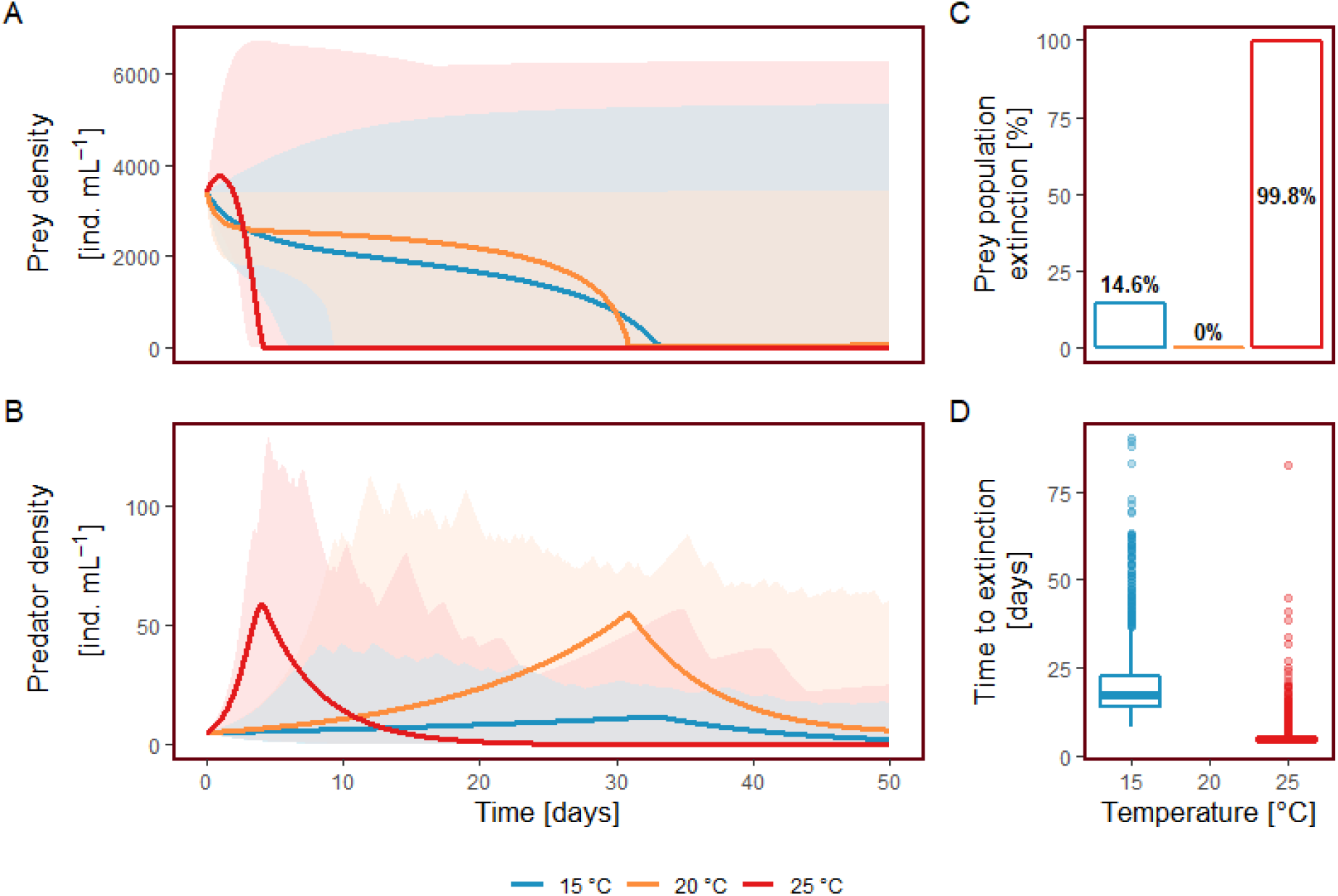
Stability analysis with the estimated model parameters and predator mortality rate set to 0.1 d^−1^, 0.156 d^−1^ and 0.241 d^−1^ at 15 °C, 20 °C and 25 °C, respectively. **A) - B)** Prey respectively predator dynamics at respective temperature. Areas denote the upper and lower trend of the simulations, while the lines represent the population dynamics with the estimated parameter values. **C)** Proportion of prey extinction in the simulations at the respective temperature. **D)** Box plots for the time to extinction in the simulations in which the prey died out at the respective temperature.

## Discussion

We show that warming can shift a predator-prey interaction from a Type III to a Type II functional response. Simulations of the dynamics of the system illustrate the ecological significance of the shift in functional response type, driving prey extinct in more than 99% of all simulations at the highest temperature, compared to less than 15% at low temperature and no extinctions at the intermediate temperature.

The change in the functional response to a Type II corroborates the findings that warming alters and potentially destabilises food webs (Petchey et al., 1999; Rall et al., 2010). However, contrary to other studies, destabilisation with warming is not caused by predator extinction through starvation (Fussmann et al., 2014; Vucic-Pestic et al., 2011) but by the prey being driven to extinction by predation. If the predator cannot switch to another prey, this will result in subsequent predator extinction. As predator-prey interactions are an integral part of food webs, such a destabilisation may have important consequences for secondary extinctions (Rall et al., 2010).

The space clearance rate exponent *q* decreased at higher temperature (with hump-shaped trend). To our knowledge, the only other study that directly investigated the link between temperature and the exponent *q* (Uszko et al., 2017) found a U-shaped trend, but the type of functional response remained consistently a Type III (*q* > 0). Elucidating the mechanisms that lead to variation in the scaling exponent *q* is a high priority for future research. In this context, Uszko et al. (2017) suggest that if the temperature approaches the thermal optimum of the predator the functional response will shift towards a Type II. Similarly, Barrios-O’Neill et al. (2016) show that in their system the functional response was closest to a Type II when the predator-prey body mass ratio was optimal for the predator. These two studies suggest that the functional response should tend towards a Type II when environmental factors converge to the respective predator optimal values.

The conversion efficiency is usually assumed to be temperature-independent in studies in which predator-prey systems are investigated under warming (Fussmann, Rosenbaum, Brose, & Rall, 2017; Uszko et al., 2017; Vasseur & McCann, 2005). However, the conversion efficiency naturally depends on the body sizes of the involved predator-prey pair and has, for instance, been calculated as 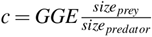 (e.g. DeLong & Luhring, 2018, *GGE* stands for gross growth efficiency, i.e. the fraction of prey biomass consumed and converted to predator biomass). Consequently, the assumption of temperature-invariant conversion efficiency might be inadequate if warming affects the predator size and prey size in a qualitatively or quantitatively different way. Indeed, the temperature-size rule (TSR, e.g. Atkinson, 1995) describes how warming reduces the body size of ectoterms, and the TSR is generally supported for protists (Atkinson, Ciotti, & Montagnes, 2003). Moreover, this effect is greater in aquatic species (Forster, Hirst, & Atkinson, 2012). Thus, under the assumption of a steeper TSR for our predator than for the smaller prey, the same amount of consumed biomass would create more new predator individuals at higher temperatures. In other words, given the right circumstances the conversion efficiency is expected to increase with warming, as observed in our study. Hence, we conclude that the conversion efficiency should not be assumed to be temperature independent *a priori*.

Temperature-dependent transitions between functional response types have previously been reported, but studies usually relied on categorical type identification using the Juliano (2001) method or similar approaches. Barrios-O’Neill et al. (2015) compared the Juliano (2001) method with a flexible approach similar to the one used in this paper and found inconsistencies in the estimated functional response types for their dataset. Consequently, the authors stated that a categorical approach to determine the functional response type such as the Juliano (2001) method may be inadequate to correctly evaluate subtle changes at low prey densities. The results of the flexible method were later confirmed by Rosenbaum and Rall (2018) when they analysed the dataset of Barrios-O’Neill et al. (2015) to test their new method. These findings raised doubts about the robustness of previously found type shifts of the functional response with warming. Our results, based on the Rosenbaum and Rall (2018) method, allowed us to confirm the presence of Type III to Type II transitions with the generalised functional response model and thus with the direct estimation of the exponent *q*.

Regardless of the method used to identify switches between functional response types, this mechanism deserves attention when the stability of predator-prey models is assessed. Besides temperature, there is mounting empirical evidence that other exposures can shift the functional response type within a predator-prey system (e.g. Barrios-O’Neill et al., 2016; Hammill, Petchey, & Anholt, 2010). Independently of whether or not the type of functional response was bounded for the estimation of the feeding rate, modelling studies that have investigated the population dynamics of predator-prey pairs across a temperature gradient limited their analysis, to our knowledge, either to a Type II (Binzer, Guill, Brose, & Rall, 2012; Fussmann et al., 2014; O’Connor, 2009; Osmond et al., 2017; Vasseur & McCann, 2005), or a Type III (Uszko et al., 2017), without including the possibility of a shifting functional response type. As switching from Type III to II represents a mechanism to destabilise consumer-resource pairs it should be included in modelling studies that assess the stability of food webs under warming, and beyond.

The iterative maximum likelihood method proposed by Rosenbaum and Rall (2018) proved to be a flexible framework to account for the particularities of our predator-system, i.e. considerable predator growth but no predator mortality. Including the numerical response without mortality rate (equation 5) led to an improved model fit even when predator growth was relatively small (in our case at 15 and 20 °C) and to a vastly improved fit when there was substantial predator growth (i.e., at 25 °C). We advise that studies that analyse the feeding rates of predator-prey systems include the predator dynamics into the estimation procedure when there is non-negligible predator growth.

The Rosenbaum and Rall (2018) method allows negative values of the space clearance rate exponent to be estimated. This exponent describes the resource dependency of the space clearance rate (i.e. *a* = *bN^q^*, see equation 1). For values of *q* below 0 the space clearance rate decreases with increasing prey density. An often chosen approach to avoid negative values for *q* is to constrain the estimation method to only fit positive values (see for instance Barrios-O’Neill et al., 2016; Van Deelen & Etter, 2003; Vucic-Pestic, Rall, Kalinkat, & Brose, 2010). Alternatively, researchers often fitted the classic functional response types (i.e. *q* = 0 for the Type II and *q* = 1 for the Type III) and thus limited the space clearance rate density dependence to predetermined categories (e.g. Seifert et al., 2014; Wollrab & Diehl, 2015). However, Rosenbaum and Rall (2018) tested their method on both published and unpublished datasets and confirmed the presence of negative values of *q* in some of them. We decided to not bound the exponent to certain ranges and indeed found it to be negative at the warmest temperature (i.e. *q* = −0.57). As this matter has received insufficient attention by the scientific community so far, hence we address it in the section “*Negative values of the space clearance rate exponent q*” and Fig. S3 in the supporting material. In the mentioned section we discuss why such a value can be biologically plausible and why the Type II functional response can be extended to encompass values of *q* greater than −1 and equal or smaller than 0. Investigating the mechanisms that can lead to a decreasing space clearance rate with increasing prey abundances represents an interesting possibility to further study predator-prey interactions.

Functional response experiments are challenging and therefore subject to logistical constraints. We were only able to test three temperature levels in our experiment. Although the temperature gradient was broad enough to detect the effect of environmental warming, a finer temperature grid (i.e. more levels) would provide more power to test the temperature scaling of the functional response parameters (Burnside et al., 2014).

Further, we decided against estimating the predator interference, considering that this would have further increased the already high complexity of the method that we applied. However, predator interference could have affected the functional response (DeLong & Vasseur, 2011). Intuitively, predator interference might cause a reduction in the space clearance rate of a predator individual, particularly with increasing predator abundance. During the 24 hours of our experiment, predator densities increased more at high prey densities, and even more so at 25 °C. Hence, predator interference could potentially explain the observed shift from a Type III to a Type II functional response at 25 °C. However, the generalised functional response (equation 1) becomes independent of the space clearance rate at high prey densities (the limit tends to the inverse of the handling time 1/*h*, i.e. the maximum feeding rate). Because of this, the distinction between functional response Type II and III (determined by the exponent *q*) depends only on the prey consumption at low prey densities. Fig. S1 in the supplementary material shows that at low prey densities the predator abundances were comparable across temperature levels. This suggests that predator interference did not influence the type of functional response. Therefore, we believe that our conclusions regarding the destabilising effect of temperature in our predator-prey system are reliable.

In conclusion, we show that warming can change the functional response of a predator by shifting it from a Type III to a Type II. The resulting increase in *per capita* prey predation at low prey abundances destabilises the predator-prey system. This finding is of considerable importance, as it represents an alternative pathway to the collapse of predator-prey systems with rising temperatures, with considerable implications for the stability of food webs. We also showed that the conversion efficiency increased with temperature and hence it should not be assumed to be temperature independent. Incorporation of these findings in future modelling work has the potential to provide more accurate predictions of community stability, structure and functioning in the face of global warming.

## Data accessibility

Data will be made available in a public repository (e.g. Dryad) upon publication.

## Acknowledgements

John P. DeLong provided valuable feedback on an earlier draft of the manuscript. Financial support for this study was granted by Swiss National Science Foundation (grant 31003A_159498 to OP) and the University of Zurich Research Priority Programme on Global Change and Biodiversity.

## Authors’ contributions

All authors conceived the ideas and designed methodology; UD and FP collected the data; UD analysed the data; UD and FP led the writing of the manuscript. All authors contributed critically to the drafts and gave final approval for publication.

