## Supplementary material for "Warming can destabilise predator-prey interactions by shifting the functional response from Type III to Type II"

#### Including predator growth and temperature-dependence of conversion efficiency

We used model selection to compare support for several biologically meaningful alternative hypotheses. First, we tested whether the inclusion of the numerical response (i.e. of the conversion efficiency $c$) improved the model fit. We compared the AIC values of the models with $c > 0$ (i.e. the unconstrained model) and without predator growth (i.e. $c = 0$). We also included a model that imposed a Type II functional response (i.e. $q = 0$) and a model where both the conversion efficiency and the exponent $q$ were set to 0 (i.e. $q = c = 0$). All comparisons were done for each temperature level separately. Model comparisons are shown in Table S1. The unconstrained model allowing for conversion efficiencies and space clearance rate exponents different from 0 consistently outperformed the other models across all temperatures (lowest AIC for the unconstrained model across temperatures).

**Table S1:** Model comparisons (AIC, log-likelihood and number of parameters). $c = 0$ means that the conversion efficiency of the predator is set to zero and thus the predator abundance is constant. With $q = 0$ the model is fixed to a Type II. The table is ordered according to decreasing ΔAIC values within temperature level.

| Temperature | Model | log-likelihood | No. of Parameters | AIC | ΔAIC |
|---|---|---|---|---|---|
| 15 °C | $q = c = 0$ | -107.20 | 5 | 224.41 | 45.92 |
| | $q = 0$ | -105.53 | 6 | 223.07 | 44.58 |
| | $c = 0$ | -83.90 | 6 | 179.81 | 1.32 |
| | **Unconstrained model** | **-82.25** | **7** | **178.49** | **0.00** |
| 20 °C | $q = 0$ | -135.77 | 6 | 283.54 | 83.26 |
| | $q = c = 0$ | -107.20 | 5 | 224.39 | 24.11 |
| | $c = 0$ | -100.34 | 6 | 212.68 | 12.40 |
| | **Unconstrained model** | **-93.14** | **7** | **200.28** | **0.00** |
| 25 °C | $c = 0$ | -233.80 | 6 | 479.61 | 162.18 |
| | $q = c = 0$ | -234.17 | 5 | 478.34 | 160.91 |
| | $q = 0$ | -160.15 | 6 | 332.31 | 14.88 |
| | **Unconstrained model** | **-151.71** | **7** | **317.43** | **0.00** |

### Parameter estimates for generalised and type II functional response across temperatures

The estimated parameter values across temperature levels are reported in Table S2. Note that the exponent $q$ of the generalised functional response was negative at 25 °C. We repeated the estimation by imposing a classical Type II functional response (i.e. $q = 0$), which allows to compare the two model fits (see Table S1) and also shows that the population stability patterns persist in both cases (see Table S4). To improve model convergence, we estimated the space clearance constant $b$ and the carrying capacity $K$ on the log-scale as they differ by several orders of magnitude.

**Table S2:** The estimates and the corresponding standard errors for the parameter values across temperatures and at 25 °C also for the model with a fixed Type II functional response. Test statistics ($z$- and $p$-values) whether the estimates are significantly different from zero are reported. The parameter $\sigma$ is the estimated standard deviation of the log-normal distributions used in the maximum likelihood procedure.

| Temperature | Type | Parameter | Estimate | Std. Error | $z$-value | $p$-value |
|---|---|---|---|---|---|---|
| 15 °C | Generalised functional response | $log(b)$ | -4.107 | 1.424 | -2.88 | 0.0039 |
| | | $h$ | 0.029 | 0.005 | 5.97 | <0.0001 |
| | | $q$ | 0.867 | 0.402 | 2.16 | 0.0309 |
| | | $r$ | 0.370 | 0.060 | 6.18 | <0.0001 |
| | | $log(K)$ | 7.902 | 0.180 | 43.98 | <0.0001 |
| | | $c$ | 0.004 | 0.002 | 1.70 | 0.0897 |
| | | $\sigma$ | 0.223 | 0.019 | 12.00 | <0.0001 |
| 20 °C | Generalised functional response | $log(b)$ | -15.230 | 0.021 | –742.35 | <0.0001 |
| | | $h$ | 0.045 | 0.005 | 9.54 | <0.0001 |
| | | $q$ | 3.493 | 0.083 | 42.14 | <0.0001 |
| | | $r$ | 1.146 | 0.067 | 17.01 | <0.0001 |
| | | $log(K)$ | 7.897 | 0.077 | 102.16 | <0.0001 |
| | | $c$ | 0.011 | 0.003 | 3.39 | 0.0007 |
| | | $\sigma$ | 0.236 | 0.020 | 12.00 | <0.0001 |
| 25 °C | Generalised functional response | $log(b)$ | 1.117 | 0.296 | 3.77 | 0.0002 |
| | | $h$ | 0.001 | 0.005 | 0.17 | 0.8653 |
| | | $q$ | -0.569 | 0.107 | -5.30 | <0.0001 |
| | | $r$ | 1.209 | 0.123 | 9.86 | <0.0001 |
| | | $log(K)$ | 8.477 | 0.165 | 51.35 | <0.0001 |
| | | $c$ | 0.010 | 0.003 | 3.58 | 0.0003 |
| | | $\sigma$ | 0.422 | 0.035 | 11.99 | <0.0001 |
| | Classic Functional response type II: $q$ fixed to 0 | $log(b)$ | -0.747 | 0.128 | -5.07 | <0.0001 |
| | | $h$ | 0.022 | 0.006 | 3.47 | 0.0005 |
| | | $q$ | 0 | n/a | n/a | n/a |
| | | $r$ | 1.269 | 0.129 | 9.86 | <0.0001 |
| | | $log(K)$ | 8.315 | 0.150 | 55.50 | <0.0001 |
| | | $c$ | 0.016 | 0.004 | 3.78 | 0.0002 |
| | | $\sigma$ | 0.426 | 0.036 | 11.72 | <0.0001 |

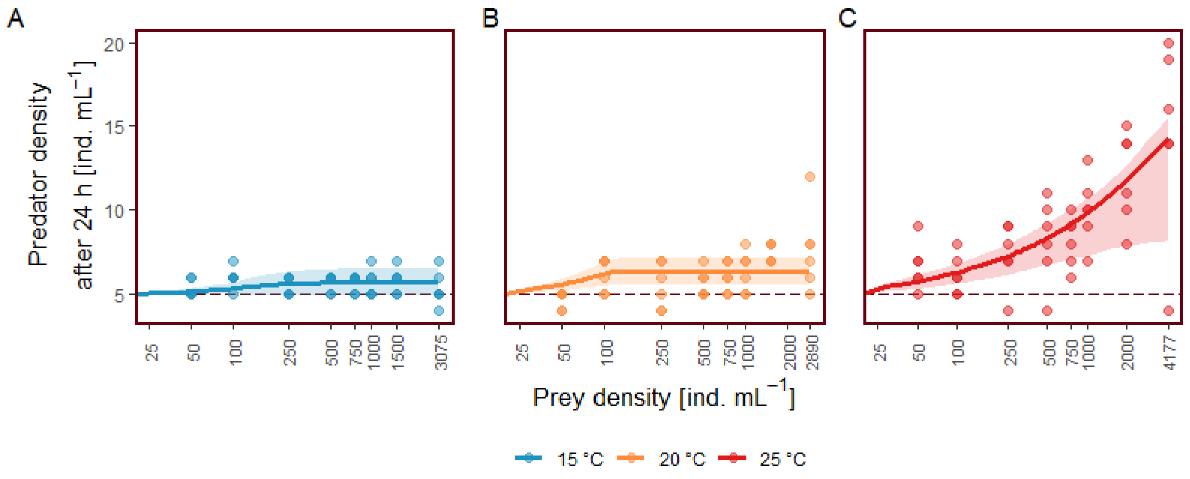


**Figure S1:** Predator dynamics. Model fits (lines) and corresponding 95% confidence intervals. Dots represent the data collected with the experiment. **A) - C)** change of predator abundance across temperature. The dashed lines represents the 5 predator individuals per millilitre present in the wells before the incubation at the start of the experiment.

### Model fit to predator dynamics

The observed predator growth during the functional response experiment was accurately captured by the model (Fig. S1). The predator growth was most pronounced at high temperatures and high prey densities.

### Estimated functional response

Fig. 1D-F in the main text displays the estimated functional responses in a linear-log plot. Fig. S2A-C shows the same plot but in a linear plot. To illustrate the differences between the estimated functional response at the different temperatures, Fig. S2D shows a zoomed in part of the estimated functional responses.

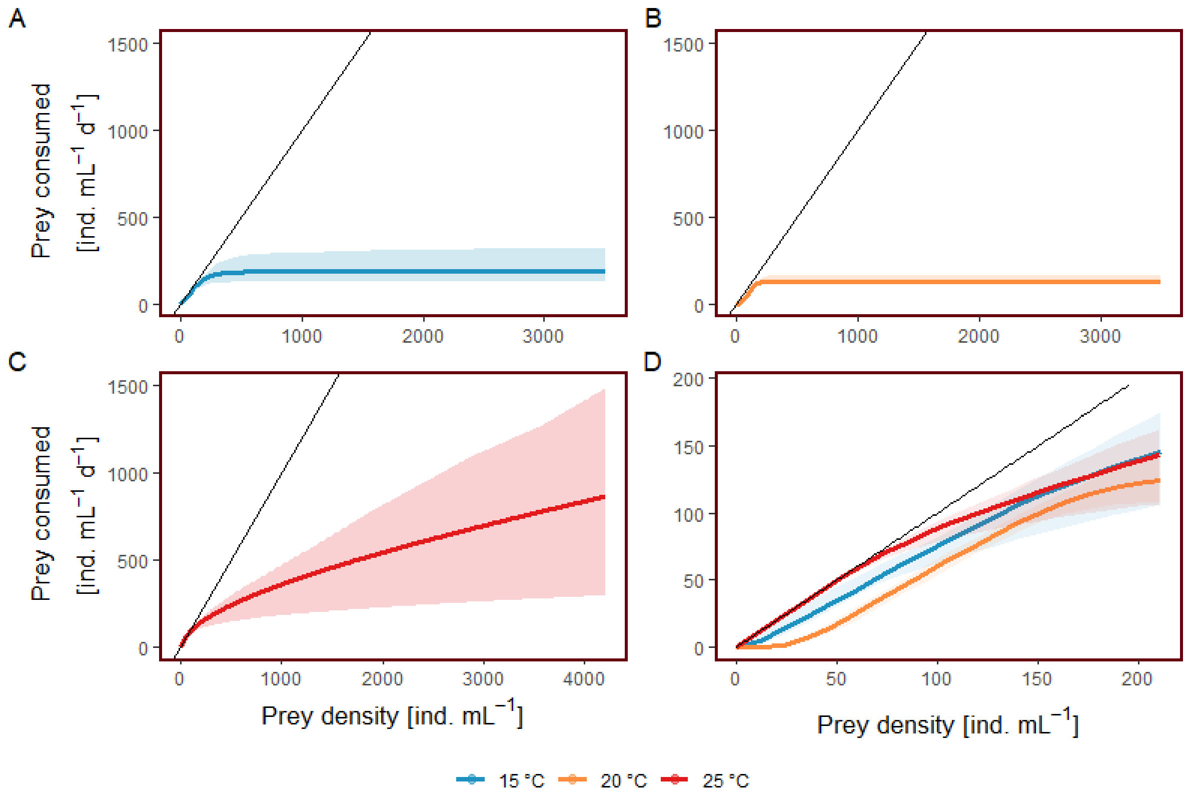


**Figure S2: A) - C)** The estimated functional response across temperature on the normal scale. For the estimation we set the prey growth rate to $r = 0$. The black line is the theoretical maximal prey consumption. **D)** A zoomed in (at low prey densities) and combined version of Fig. S2A-C.

### Estimated temperature and prey density dependency of the space clearance rate

The space clearance rate is modelled with the equation $a = bN^q$, hence both the exponent $q$ and the constant $b$ are needed for its interpretation (Fig. S3). The space clearance rate increased with prey density at the temperatures 15 and 20 °C. Due to the negative value of the attack exponent $q$ at 25 °C ($q_25 = -0.57$, see Table S2) the space clearance is estimated to decrease with increasing prey abundance at the highest temperature. Fixing the functional response to a Type II, means fixing the exponent $q$ to 0. In this case the space clearance rate is independent of the resource (i.e. the prey abundance) and equal to estimated space clearance constant $b$.

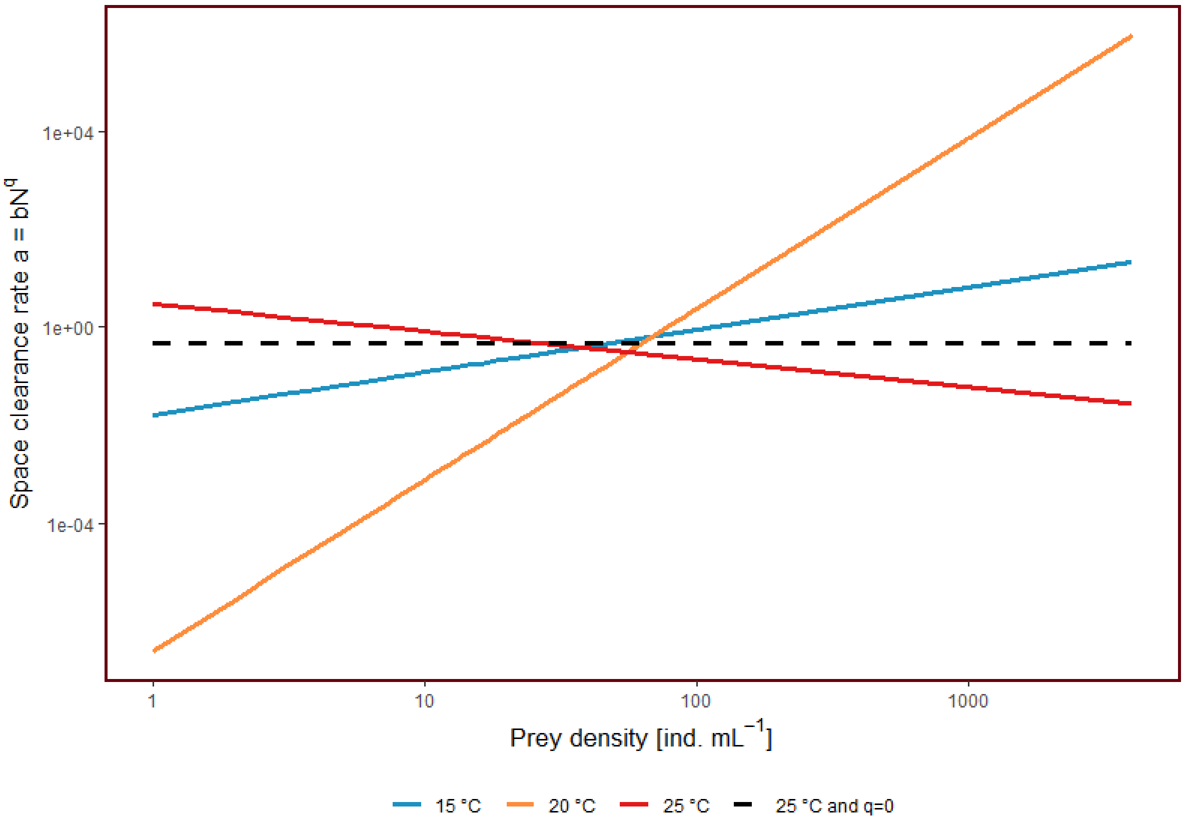


**Figure S3:** The estimated space clearance rate $a = bN^q$ across temperature on the log10-log10 scale. Parameter values are reported in Table S2. Also included is the model at 25 °C for which the functional response was fixed to a Type II ($q = 0$).

### Sensitivity analysis of predator mortality

Predator mortality was not measured experimentally. We chose it based on the procedure described in the subsection "*Stability analysis*" in the methods section of the main text. To assess the robustness of our results, we tested population stability across a range of predator mortality rates (Table S3).

Table S4 shows the results of the stability analysis across different predator mortality rates (based on Table S3) and for different models: the simulations were repeated for the model with flexible exponent $q$ and for the model with fixed $r$, $K$, $c$ and $\sigma$ values (fixed at the values estimated at 20 °C). We drew the parameter combinations from a multivariate normal distribution (mean =

**Table S3:** Mortality rates for the predator across temperatures.

| Simulation | Predator mortality rate | | |
|---|---|---|---|
| | at 15 °C | at 20 °C | at 25 °C |
| 1 | 0.100 $d^{-1}$ | 0.156 $d^{-1}$ | 0.241 $d^{-1}$ |
| 2 | 0.200 $d^{-1}$ | 0.313 $d^{-1}$ | 0.481 $d^{-1}$ |
| 3 | 0.300 $d^{-1}$ | 0.469 $d^{-1}$ | 0.722 $d^{-1}$ |

estimated parameter values, variance = covariance matrix of model fit). The respective correlation matrices are shown in Table S5. For the model with fixed $r$, $K$, $c$ and $\sigma$ values we restricted the covariance matrix to only contain the non-fixed parameters. At 25 °C, we carried out the sensitivity analysis also for the model for which we imposed a classic Type II functional response (i.e. $q = 0$) in the estimation. Please note that in some of the simulations, no extinctions happened (Table S4).

The sensitivity analysis showed that the patterns persisted (although attenuated) across all mortality rates and for all tested models: at 25 °C the predator drove the prey extinct decisively more often than at the other two temperatures. In fact, at 25 °C the prey went extinct in more simulations and also faster than at 15 and 20 °C. This illustrates the stabilising properties of the Type III functional response compared to the Type II.

### Negative values of the space clearance rate exponent $q$

In this study, we did not bound the exponent $q$ to certain ranges or values, thus allowing it to take on negative values. We estimated the exponent to be negative at the warmest temperature (i.e. $q_{25} = -0.57$). Here we explore whether such a result is reasonable or not, starting with the definition of the space clearance rate.

For his famous disc equations, Holling (1959) introduced the space clearance rate calling it the instantaneous rate of discovery and defined it as the search rate multiplied with the detection probability (unit: $m^2\,s^{-1}$ or $m^3\,s^{-1}$). Additionally to the search rate and the detection probability, the space clearance rate is nowadays considered to represent even more underlying processes oc-

**Table S4:** Percentage of prey persistence / extinction and the mean time to extinction (–, if no extinctions happened) and its standard deviation for simulations of community dynamics across temperatures. The simulations were repeated for the model with flexible exponent $q$, for the model with fixed $r$, $K$, $c$ and $\sigma$ values and at 25 °C also for the Type II model (i.e. $q = 0$).

| Temper-ature | Model type | Predator mortality rate | Simulations with prey persistence | Simulations with prey extinction | Mean time to prey extinction | Standard deviation |
|---|---|---|---|---|---|---|
| 15 °C | Flexible | $0.100\,d^{-1}$ | 85.4 % | 14.6 % | 20.4 d | 10.2 d |
| | | $0.200\,d^{-1}$ | 98.4 % | 1.6 % | 26.2 d | 13.5 d |
| | | $0.300\,d^{-1}$ | 100.0 % | 0.0 % | 45.0 d | 17.6 d |
| | Fixed $r$, $K$, $c$ and $\sigma$ | $0.100\,d^{-1}$ | 64.9 % | 35.1 % | 8.8 d | 1.9 d |
| | | $0.200\,d^{-1}$ | 75.4 % | 24.6 % | 11.8 d | 3.5 d |
| | | $0.300\,d^{-1}$ | 85.4 % | 14.6 % | 18.5 d | 10.3 d |
| 20 °C | Flexible | $0.156\,d^{-1}$ | 100.0 % | 0.0 % | - | - |
| | | $0.313\,d^{-1}$ | 100.0 % | 0.0 % | - | - |
| | | $0.469\,d^{-1}$ | 100.0 % | 0.0 % | - | - |
| | Fixed $r$, $K$, $c$ and $\sigma$ | $0.156\,d^{-1}$ | 100.0 % | 0.0 % | - | - |
| | | $0.313\,d^{-1}$ | 100.0 % | 0.0 % | - | - |
| | | $0.469\,d^{-1}$ | 100.0 % | 0.0 % | - | - |
| 25 °C | Flexible | $0.241\,d^{-1}$ | 0.2 % | 99.8 % | 5.1 d | 2.1 d |
| | | $0.481\,d^{-1}$ | 4.3 % | 95.7 % | 8.0 d | 4.4 d |
| | | $0.722\,d^{-1}$ | 70.9 % | 29.1 % | 12.8 d | 4.7 d |
| | Fixed $r$, $K$, $c$ and $\sigma$ | $0.241\,d^{-1}$ | 0.0 % | 100.0 % | 5.0 d | 1.7 d |
| | | $0.481\,d^{-1}$ | 13.9 % | 86.1 % | 8.5 d | 4.4 d |
| | | $0.722\,d^{-1}$ | 92.1 % | 7.9 % | 6.2 d | 1.8 d |
| | Fixed to Type II | $0.241\,d^{-1}$ | 0.0 % | 100.0 % | 6.3 d | 1.0 d |
| | | $0.481\,d^{-1}$ | 0.3 % | 99.7 % | 11.7 d | 3.3 d |
| | | $0.722\,d^{-1}$ | 67.6 % | 32.4 % | 41.6 d | 23.0 d |

curring in a natural predator-prey system. Such processes are for instance the success probability of attacks, the encounter rate and the attack efficiency (e.g. Jeschke, Kopp, & Tollrian, 2002).

Every process implicitly modelled with the space clearance rate can potentially depend on a number of other attributes of the predator-prey system. For instance, Vucic-Pestic, Ehnes, Rall, and Brose (2011) argue that the encounter rate depends on the movement speed of both the prey and the predator, while the success rate is inversely proportional on the preys escape efficiency. Hence, many mechanisms could potentially cause the space clearance rate to decrease with increasing resource abundance.

Following the generalised functional response equation (equation 1 in the main text) the functional response type, and hence the value of $q$, is determined by the response to low prey densities. As we explain in the main text, this combined with the comparable predator abundances at low prey densities makes predator inference unlikely to be the cause of the decreasing space clearance rate in this model framework. Similarly, predator satiation, which suggests a decrease in space clearance rate because of low hunger levels at high prey density, cannot explain the functional response type shift. However, density-dependent behaviour of the prey, such as increased behavioural defence caused by the prey occasionally forming ”swarms” (personal observation) could potentially explain the decreasing space clearance rate with increasing prey abundance.

The plausibility of a negative exponent is confirmed mathematically: for $q > -1$ the maximum feeding rate (prey density tending to infinity in equation 1) tends to $1/h$ (i.e. the inverse of the handling time), while for $q < -1$ it tends to 0 (which is biologically implausible). For the special case of $q = -1$, equation 1 reflects the unlikely scenario of feeding rates that are independent of prey abundance. Moreover, for $-1 < q < 0$ the functional response retains the form of a Type II (i.e. a hyperbolic curve, see Fig. S1).

For these reasons, we suggest that the type II definition should be broadened from $q = 0$ to $-1 < q \leq 0$ and thus to include space clearance rates that range from having an inverse relation with prey abundance to being resource independent. The Type I and Type III definitions remain unchanged.

### Interdependence of model parameters

Table S5 displays the correlation matrices of the model fit at the three temperatures. As can be seen, the parameters $b$ and $q$ are strongly correlated across all temperatures. This is expected as they are the two parameters that determine the space clearance rate and hence also the shape of the functional response. Similarly, and also as expected as it determines the functional response as well,

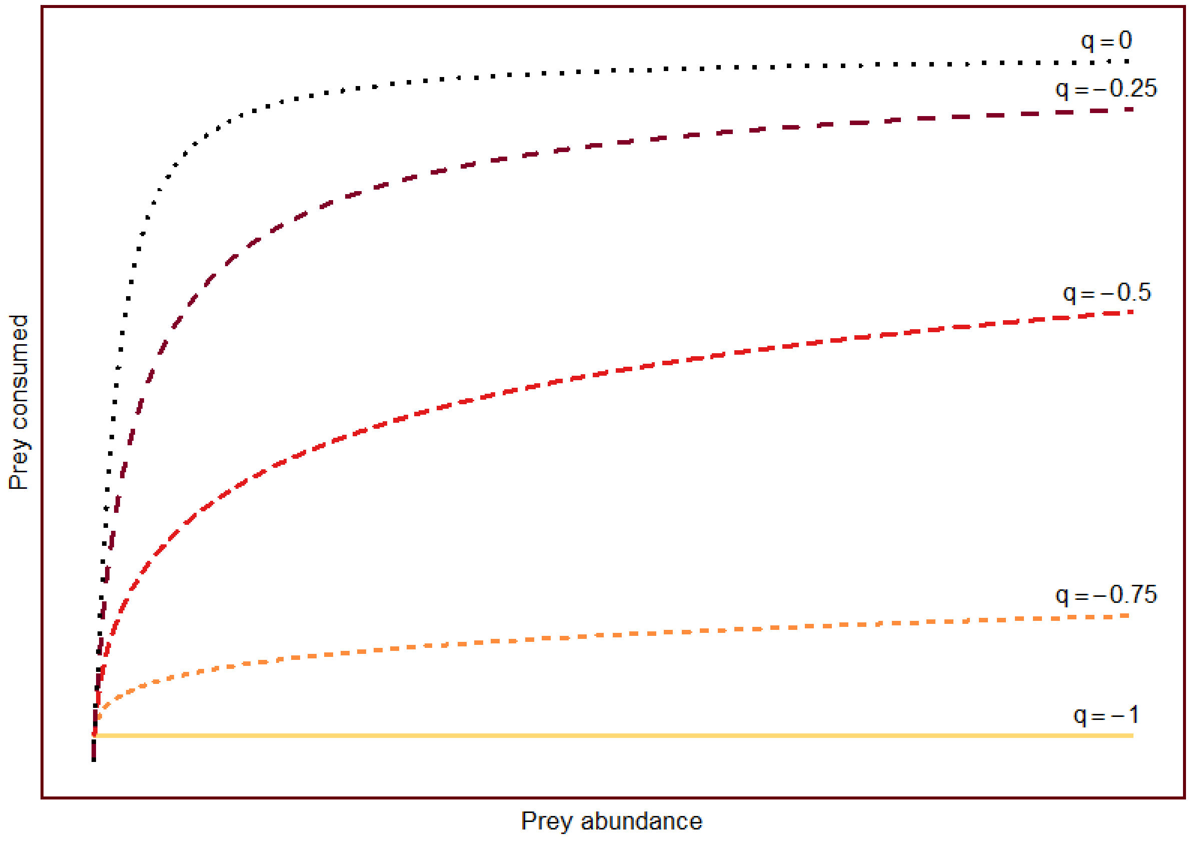


**Figure S4:** Functional response shape ranging from a classical Type II ($q = 0$) to a resource independent response ($q = -1$). For $q$ values greater than -1 and smaller or equal to 0 the functional response is a hyperbolic curve.

the handling time correlates with space clearance rate parameters as well, but less so. The other model parameters do not show any strong correlations that would compromise their independent interpretation and the validity of the results.

### Model improvement

The Rosenbaum and Rall (2018) method including intrinsic prey growth and prey carrying capacity and combined with the predator dynamics successfully captured the characteristics of our predator-prey system. Nevertheless, improvements are still possible and other predator-prey systems will have other characteristics that will need to be accounted for. For instance, we were able to ignore the predator mortality parameter in the estimation procedure, as it was negligible in our experimental setup. Should this not be the case, one can estimate the predator mortality rate in the same way as the conversion efficiency, i.e. by using equation 4 instead of equation 5 for the model estimation (see main text). However, this could prove to be challenging and unsuccessful as the mortality rate $m$ and the conversion efficiency $c$ are likely to be strongly correlated. Alternatively, the predator mortality rate can be estimated with an independent experiment or be based on published mortality estimates. The predator mortality rate can then be fixed in equation 4 to obtain an independent estimate of the conversion efficiency.

As the estimation method that we used in our study is based on time series, the method can most likely be improved by measuring the prey (and predator) abundance at intermediate time steps. Whether this is possible, however, depends on the characteristics of the setup, namely on whether this measurement can be taken without compromising the experimental units. In our case, this was not possible because we used the medium of each experimental well to estimate abundances (see method section in the main text). If intermediate abundance counts are to be taken, the Rosenbaum and Rall (2018) estimation method needs to be changed accordingly.

Finally, in functional response experiments the prey abundance gradient is of crucial importance: it strongly influences the reliability of the parameter estimates. Particularly, high prey abundances are needed to estimate the handling time. Similarly, low prey abundance levels are needed to estimate the shape and the steepness of the functional response (i.e. the space clearance parameters).

**Table S5:** Correlation matrices for the model fits at the temperatures 15, 20 and 25 °C

Correlation matrix of the model fit at 15 °C

| | *b.log* | *h* | *q* | *r* | *K.log* | *c* | $\sigma$ |
|---|---|---|---|---|---|---|---|
| *b.log* | 1 | | | | | | |
| *h* | -0.707 | 1 | | | | | |
| *q* | -0.997 | 0.720 | 1 | | | | |
| *r* | 0.324 | -0.658 | -0.311 | 1 | | | |
| *K.log* | 0.103 | -0.064 | -0.116 | -0.256 | 1 | | |
| *c* | -0.159 | 0.441 | 0.159 | -0.177 | -0.011 | 1 | |
| $\sigma$ | -0.004 | 0.005 | 0.004 | -0.003 | -0.001 | 0.001 | 1 |

Correlation matrix of the model fit at 20 °C

| | *b.log* | *h* | *q* | *r* | *K.log* | *c* | $\sigma$ |
|---|---|---|---|---|---|---|---|
| *b.log* | 1 | | | | | | |
| *h* | 0.349 | 1 | | | | | |
| *q* | 1.000 | 0.345 | 1 | | | | |
| *r* | 0.109 | -0.641 | 0.115 | 1 | | | |
| *K.log* | -0.116 | 0.175 | -0.117 | -0.439 | 1 | | |
| *c* | 0.069 | 0.553 | 0.068 | -0.226 | 0.070 | 1 | |
| $\sigma$ | -0.008 | -0.007 | -0.008 | 0.000 | 0.001 | 0.002 | 1 |

Correlation matrix of the model fit at 25 °C

| | *b.log* | *h* | *q* | *r* | *K.log* | *c* | $\sigma$ |
|---|---|---|---|---|---|---|---|
| *b.log* | 1 | | | | | | |
| *h* | -0.783 | 1 | | | | | |
| *q* | -0.975 | 0.833 | 1 | | | | |
| *r* | -0.381 | 0.294 | 0.528 | 1 | | | |
| *K.log* | -0.110 | -0.073 | 0.032 | -0.303 | 1 | | |
| *c* | 0.452 | 0.060 | -0.470 | -0.596 | -0.137 | 1 | |
| $\sigma$ | 0.033 | -0.047 | -0.037 | -0.019 | 0.012 | -0.003 | 1 |
